# Abl2:cortactin interactions regulate dendritic spine stability via control of a stable filamentous actin pool

**DOI:** 10.1101/2020.09.12.294694

**Authors:** Juliana E. Shaw, Michaela B. C. Kilander, Yu-Chih Lin, Anthony J. Koleske

## Abstract

Dendritic spines are enriched with stable and dynamic actin filaments, which determine their structure and shape. Disruption of the Abl2/Arg nonreceptor tyrosine kinase in mice compromises spine stability and size. We provide evidence that binding to cortactin tethers Abl2 in spines, where Abl2 and cortactin maintain the small pool of stable actin required for dendritic spine stability. Using fluorescence recovery after photobleaching of GFP-actin, we find that disruption of Abl2:cortactin interactions eliminates stable actin filaments in dendritic spines, significantly reducing spine density. A subset of spines remaining after Abl2 depletion retain their stable actin pool and undergo activity-dependent spine enlargement associated with increased cortactin levels. Finally, tonic increases in synaptic activity rescue spine loss upon Abl2 depletion by promoting cortactin enrichment in vulnerable spines. Together, our findings strongly suggest Abl2:cortactin interactions promote spine stability by maintaining pools of stable actin filaments in spines.

## Introduction

Dendritic spines are protrusions that extend from dendrites and serve as receptive post-synaptic compartments on excitatory neurons. Defects in dendritic spine formation, spine density and spine shape are hallmarks of many brain disorders (Blanpied & Ehlers, 2004; Fiala, Spacek, & Harris, 2002; Herms & Dorostkar, 2016; Hu, Shih, Shih, & Hsueh, 2016; Y. C. Lin & Koleske, 2010; Penzes, Cahill, Jones, VanLeeuwen, & Woolfrey, 2011; Spence & Soderling, 2015; van Spronsen & Hoogenraad, 2010). Spine shape and stability are principally supported by a filamentous actin network, which also organizes scaffolding proteins and neurotransmitter receptors at the post- synaptic density (Alvarez & Sabatini, 2007; Fischer, Kaech, Knutti, & Matus, 1998; Hotulainen & Hoogenraad, 2010; W. H. Lin & Webb, 2009; Schubert & Dotti, 2007). Spine actin filaments are assembled into higher-order structures that undergo dynamic rearrangements and provide the forces to drive changes in spine shape and size (Chazeau & Giannone, 2016; Luo, 2002; Matus, 2000; Matus, Brinkhaus, & Wagner, 2000; Sekino, Kojima, & Shirao, 2007). Indeed, molecules that regulate spine plasticity and stability, including cell surface receptors and kinases, often converge on signaling pathways that control the polymerization, stability, and contractility of actin networks. Therefore, the cytoskeletal machinery is a key module to regulate spine plasticity and stability.

Fluorescence recovery after photobleaching (FRAP) experiments of GFP-actin have determined that most (80-85%) spine actin filaments are dynamic, undergoing ongoing polymerization and turnover within 10s of seconds, while the remainder of actin (10-20%) is more stable. These distinct kinetic pools of actin likely play unique roles in the spine. Dynamic filaments mediate a net retrograde flow of actin from the spine periphery toward the spine core and are perfectly positioned to promote changes in spine size and shape (Frost, Shroff, Kong, Betzig, & Blanpied, 2010; Honkura, Matsuzaki, Noguchi, Ellis-Davies, & Kasai, 2008) and also regulate neurotransmitter receptor localization or gating properties (Borgdorff & Choquet, 2002; Kerr & Blanpied, 2012; Rosenmund & Westbrook, 1993). Photoactivation of mEOS3.2-actin reveals that the small stable pool of actin is enriched at the spine base and turns over much more slowly (t ~17 minutes)(Honkura et al., 2008). The relative proportion of actin in this stable pool increases during synapse maturation (Koskinen, Bertling, Hotulainen, Tanhuanpaa, & Hotulainen, 2014). This stable pool may serve as a base for polymerization of new actin during spine enlargement (Mikhaylova et al., 2018). The functions of the stable actin pool have been largely understudied, primarily because we lack an in-depth understanding of the molecules and mechanisms that regulate it.

The Abl2/Arg nonreceptor tyrosine kinase interacts with its substrate and binding partner cortactin to regulate actin-based structures in many cellular contexts (MacGrath & Koleske, 2012b; Schnoor, Stradal, & Rottner, 2018). Both proteins localize to dendritic spines where they promote spine stability (Hering & Sheng, 2003; Iki, Inoue, Bito, & Okabe, 2005; Y. C. Lin, Yeckel, & Koleske, 2013; MacGillavry, Kerr, Kassner, Frost, & Blanpied, 2016; Mikhaylova et al., 2018; Sfakianos et al., 2007). Abl2 binds actin filaments cooperatively (Wang, Miller, Mooseker, & Koleske, 2001) and increases the binding stoichiometry of cortactin for actin filaments (MacGrath & Koleske, 2012a). These interactions significantly impact actin filament stability – at saturated binding, Abl2 and cortactin each stabilize actin filaments (Courtemanche, Gifford, Simpson, Pollard, & Koleske, 2015; Scherer, Anand, & Koleske, 2018), but mixtures of Abl2 and cortactin at concentrations far below saturated binding also stabilize actin filaments (Courtemanche et al., 2015). While Abl2 and cortactin synergize to stabilize actin filaments *in vitro*, it is unclear whether or how they contribute to actin stability in neurons or how this mechanism might impact dendritic spine shape or stability.

Here, we provide evidence that cortactin anchors Abl2 in dendritic spines via an SH2:phosphotyrosine interface, and mutational disruption of this interface dislodges Abl2 from spines. Depletion of either protein eliminates the pool of stable actin in spines, as measured by FRAP of GFP-actin. Disruption of this stable pool correlates with a significant loss of dendritic spines. We find that a subset of spines remaining after Abl2 depletion exhibit increased spine head widths, which is tightly associated with increased cortactin levels and retention of the stable actin pool. Finally, tonic increases in activity can rescue spine loss upon Abl2 depletion by promoting cortactin enrichment in vulnerable spines that are otherwise lost in basal conditions. Together, our data indicate that Abl2 and cortactin synergize to maintain dendritic spine stable actin, which is critical for spine stability.

## Results

### Abl2 kinase activity and SH2 domains are critical for Abl2 enrichment in dendritic spines

Abl2 is a large multidomain protein that contains N-terminal Src homology (SH) 3, SH2, and kinase domains, followed by a C-terminal extension that contains a proline rich PXXP segment, and two F-actin binding domains flanking a microtubule-binding domain (Figure 1A). When expressed in wild type neurons, Abl2-RFP preferentially localizes to spines. While Abl2 KD decreases spine density, overexpression does not alter spine density (Y. C. Lin et al., 2013). To study the relative contributions of different Abl2 protein domains to its spine enrichment, we expressed different RFP-tagged Abl2 point and truncation mutants (Figure 1A) along with GFP as a cell fill (Figure 1B,C). Consistent with previous studies (Y. C. Lin et al., 2013), we find Abl2-RFP is enriched approximately 2-fold in spines relative to dendritic shafts in control neurons (Figure 1B,C). The Abl2 N-terminal half (Abl2 N-term-RFP), which contains the SH3, SH2, and kinase domains was enriched in spines to a similar extent as full-length Abl2, while the Abl2 C-terminal half (Abl2 C-term-RFP) was not (Figure 1B,C). A kinase-inactive point mutant of Abl2 (Abl2-KI-YFP) exhibited significantly less enrichment (Figure 1C), supporting a key role for kinase activity in proper localization of Abl2 to dendritic spines. We also mutated the Abl2 SH2 domain (R198K) to abrogate binding to phospho-tyrosine-containing binding partners (Lapetina, Mader, Machida, Mayer, & Koleske, 2009; Simpson et al., 2015), both in full length Abl2 and in Abl2 N-term (Figure 1A). While both mutants were present in spines, neither was enriched relative to the dendrite shaft (Figure 1B,C). Together, these data indicate that Abl2 requires both its kinase activity and SH2 domains to support its preferential localization to dendritic spines.

**Figure 1.**
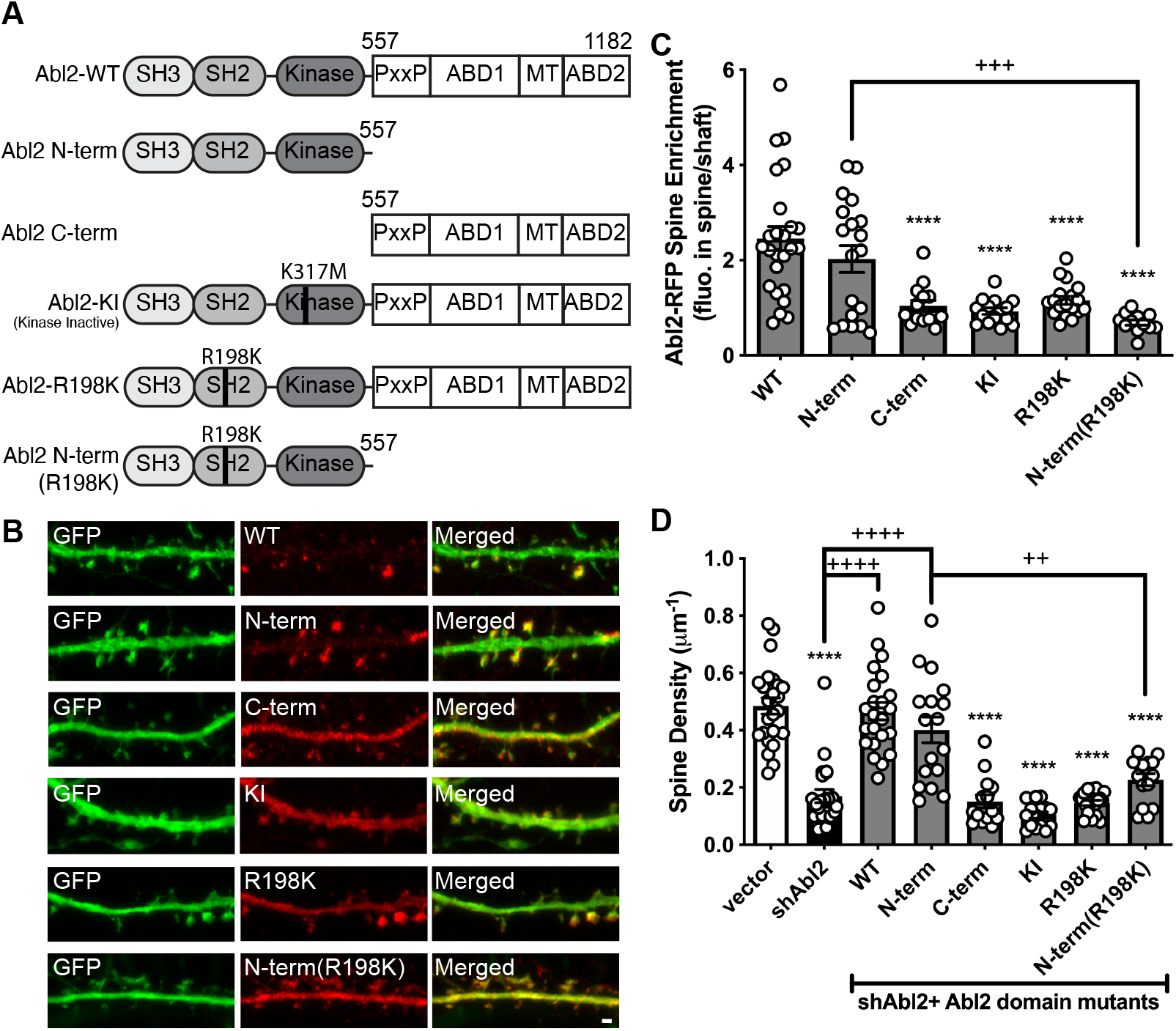
Abl2 kinase and SH2 domain are required for spine localization and spine stability. **A.** Illustration of Abl2 domain mutants used in the experiment. **B.** Representative confocal images of neurons co-transfected with GFP and RFP-tagged Abl2 or Abl2 domain. Scale bar = 1 μm. **C.** The relative fluorescence intensity (spine enrichment) of Abl2-RFP or Abl2 domain mutants in the dendritic spine vs dendrite shaft. Note that spine enrichment > 1 indicates a preferential localization to dendritic spines. Data are means + SEM (n = 12-25 neurons/ group). +++p < 0.001, ****p < 0.0001 (ordinary one-way ANOVA with Tukey multiple comparisons test). **D.** Dendritic spine density of neurons transfected with empty vector control, shAbl2, or shAbl2 with Abl2/Abl2 domain mutants. Note only neurons transfected with Abl2 domain mutants that contain a functional kinase domain and a SH2 domain (e.g. Abl2-WT and Abl2 N-term) show spine density comparable to control neurons. Data are means + SEM (n = 14-24 neurons/ group). *p < 0.05, ++p < 0.01, ****,++++p < 0.0001 (ordinary one-way ANOVA with Tukey multiple comparisons test).

### Abl2 kinase activity and SH2 domains are required for dendritic spine maintenance

We previously showed that knockdown of Abl2 in hippocampal neurons caused progressive reduction of dendritic spines over 72 hours and enlargement of a subset of remaining spines (Y. C. Lin et al., 2013). These defects were rescued by re-expression of the shRNA-resistant Abl2. We used this complementation assay to determine which domains of Abl2 are important to maintain normal dendritic spine density (Figure 1A). Similar to our previous work (Y. C. Lin et al., 2013), we found that Abl2 KD yielded a 65 ± 8% decrease in spine density after 72 hours and the full-length shRNA-resistant Abl2-YFP restored spine density to levels observed in the vector alone control. Re-expression of the kinase-inactive (Abl2-KI-YFP) and SH2 domain mutant (Abl2-R198K-YFP) failed to rescue spine density (Figure 1D), which paralleled their lack of enrichment in dendritic spines (Figure 1C). Surprisingly, the Abl2 N-term-YFP, which completely lacks the C-terminal actin binding domains, was sufficient to support normal dendritic spine density, while the Abl2 C-terminal half (Abl2-C-term-YFP) did not (Figure 1D). Abl2 constructs that are enriched in dendritic spines (Figure 1C) support spine densities (Figure 1D) similar to WT Abl2, while mutants that disrupt spine enrichment do not support normal spine density (summarized in Table 1). Hence, Abl2 localization to dendritic spines is critical for spine stability.

**Table 1.** Summary of Abl2 and cortactin spine enrichment and spine maintenance.

|  |  | **Spine**<br>**Enrichment** | **Normal spine**<br>**density?** |  |
| --- | --- | --- | --- | --- |
|  |  |  |  | **Cortactin spine**<br>**enrichment** |
| **Abl2** |  | + | Y |  |
|  | Abl2 KD | - | N | reduced |
|  | Abl2 N-term | + | Y | normal |
|  | Abl2 C-term | - | N |  |
|  | Abl2-KI | - | N |  |
|  | Abl2-R198K | - | N |  |
|  | Abl2 N-term<br>(R198K) | - | N |  |
|  |  |  |  | **Abl2 spine**<br>**enrichment** |
| **cortactin** |  | + | Y |  |
|  | Cort KD | - | N | reduced |
|  | 3YFcort | + | N | reduced and<br>shows increased<br>turnover |
|  | W525Acort | - | N |  |

+: nearly WT spine enrichment of the mutant; -: no preferential enrichment in the spine compared to the dendrite shaft.

### Cortactin tethers Abl2 in dendritic spines to support normal dendritic spine density

Abl2 kinase activity and phosphotyrosine-binding by its SH2 domain are necessary for Abl2 to both localize to dendritic spines and support normal dendritic spine density, but the critical partners for Abl2 in these processes are unknown. Abl2 and cortactin co-localize to actin rich protrusions in a variety of cellular contexts and reciprocal interactions between the two proteins have been well characterized (Courtemanche et al., 2015; Gifford et al., 2014; Y. C. Lin et al., 2013; Liu, MacGrath, Koleske, & Boggon, 2012). The cortactin SH3 domain binds to a specific Pro-X-X-Pro-X-X-Pro motif unique to Abl2 (Lapetina et al., 2009; Liu et al., 2012). Subsequent Abl2-mediated phosphorylation of cortactin on tyrosine (Y)421 and Y466 creates binding sites for the Abl2 SH2 domain (Gifford et al., 2014).

We asked whether Abl2 enrichment in spines depends on cortactin, and if so, whether interactions between the proteins are critical for their enrichment. We designed two shRNAs predicted to target cortactin: shcort #1 and shcort #2. Transfection of shcort #1 in HEK-293 cells resulted in a 74 ± 9% (difference of means ± SEM) reduction of a mouse cortactin-RFP fusion protein. Shcort #2 did not impact cortactin-RFP expression (Figure 2A,C), therefore we used shcort #1 (shcort) to knock down cortactin in subsequent experiments. Levels of an shRNA-resistant cortactin (cort^r^-RFP), bearing silent mutations that disrupted interaction with shcort #1, were unchanged up to 72 hrs after transfection with shcort #1 (Figure 2B,C).

**Figure 2.**
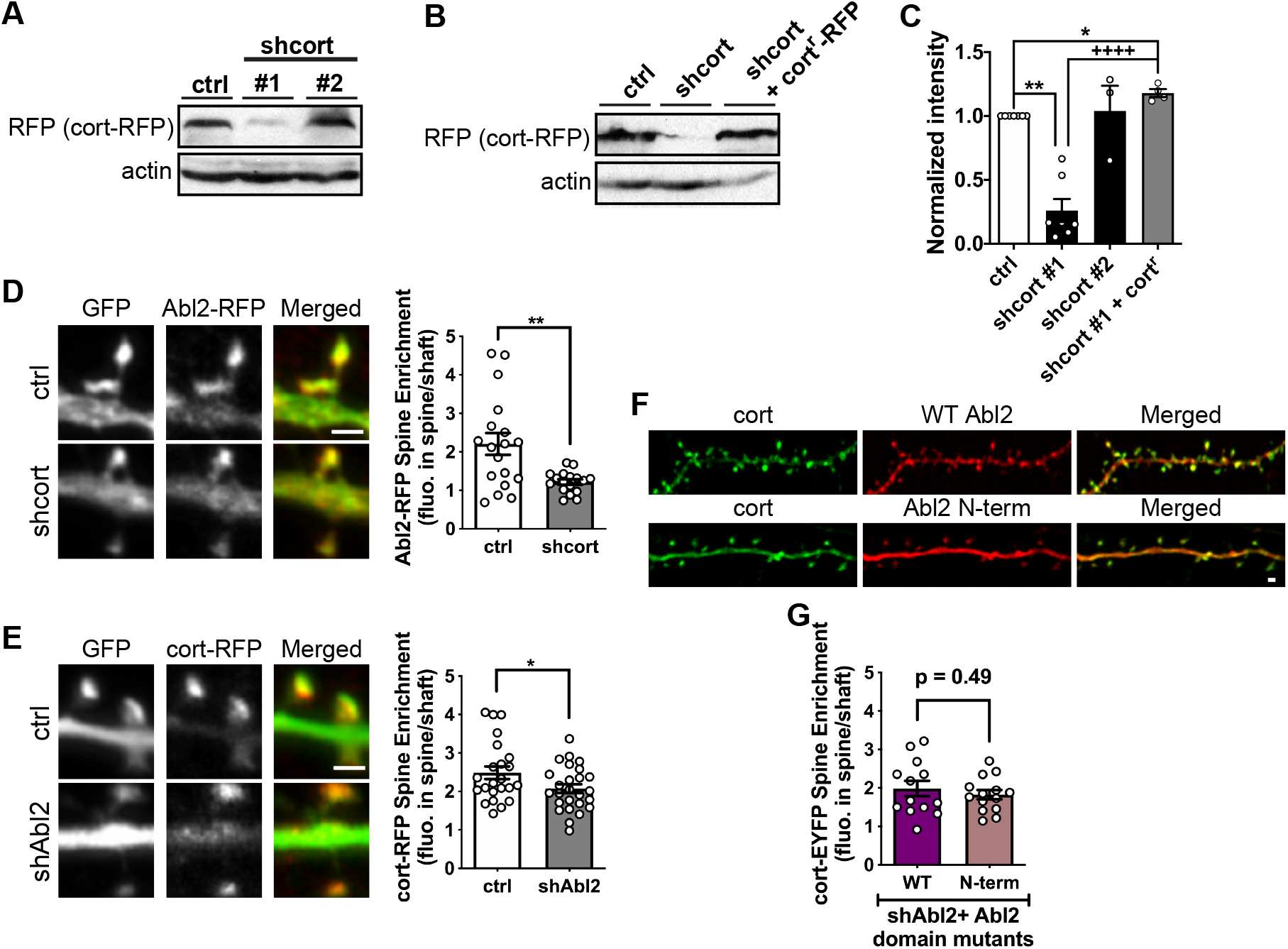
Abl2 enrichment in dendritic spines is strongly dependent on cortactin levels. **A.** Representative immunoblots of cortactin KD in HEK-293 cells. Two shRNA constructs (shcort #1 and shcort#2) were co-transfected with cortactin-RFP in HEK293 cells for 72 hours. The expression of cortactin-RFP was determined using RFP antibodies on Western blot analysis. Shcort #1 (later as shcort) has higher KD efficiency than shcort #2. **B.** HEK-293 cells were co-transfected with control vector or shcort together with cortactin-RFP or cort^r^-RFP, an shRNA resistant form of cortactin, for 72 hours. Shcort efficiently knocks down cortactin-RFP, but not cort^r^-RFP. **C.** Quantification of cortactin-RFP levels indicating shRNA KD of cortactin and resistance of cort^r^-RFP towards the shRNA. Data are means + SEM (n = 3-7 cultures/condition) normalized to the actin signal. ****,++++p < 0.0001 (one sample t test). **D.** Representative confocal images and relative enrichment of Abl2-RFP in cortactin KD neurons. Scale bar = 1 μm. When cortactin is knocked down, Abl2 shows less spine enrichment. Data are means + SEM (n = 17-18 neurons/group). **p < 0.01 (unpaired t test). **E.** Representative confocal images and relative enrichment of cort-RFP in Abl2 KD neurons. Scale bar = 1 μm. Data are means + SEM (n = 23-26 neurons/group). *p < 0.05 (unpaired t test). **F.** Representative confocal images of neurons co-transfected with cortactin-EYFP and RFP-tagged Abl2 or Abl2 N-term in Abl2 KD neurons. Scale bar = 1 μm. **G.** The relative enrichment of cort-EYFP in dendritic spines of neurons expressing Abl2 or Abl2 N-term. Cortactin enrichment in spines is comparable in each condition. Data are means + SEM (n = 13-14 neurons/group). p = 0.49 (unpaired t test).

We knocked down Abl2 or cortactin in cultured hippocampal neurons to examine how these manipulations impacted the subcellular localization of its reciprocal binding partner. Consistent with Figure 1C and previous studies, we find Abl2-RFP and cort-RFP are enriched approximately 2-fold in dendritic spines relative to shafts in control neurons (Figure 2D,E). The enrichment of Abl2-RFP in dendritic spines relative to the dendritic shaft was decreased by 45 ± 14% (Figure 2D) in cortactin KD neurons, demonstrating that spine enrichment of Abl2 is strongly dependent on the presence of cortactin. These data indicate that cortactin is an important tether for Abl2 in dendritic spines. In contrast, the relative enrichment of cortactin-RFP in dendritic spines was reduced by 17 ± 8% in Abl2 KD neurons (Figure 2E)(Y. C. Lin et al., 2013). Abl2 KD neurons re-expressing full length Abl2 or Abl2 N-term also exhibited a 2-fold enrichment of cort-EYFP in spines (Figure 2F,G), an enrichment similar to that observed in control neurons (Figure 2E), indicating that cortactin enrichment in spines only partially depends on Abl2.

We next examined the critical determinants in cortactin required for its enrichment in spines and its ability to support Abl2 enrichment in spines. We first measured the enrichment of shRNA resistant cortactin-RFP and mutant constructs in neurons following treatment with shcort (Figure 3A-C). Cortactin bearing non-phosphorylatable tyrosine-to-phenylalanine substitutions at Y421, Y466, and Y482 (3YFcort^r^-RFP, Figure 3A) was enriched in spines more than WT cort^r^-RFP (Figure 3B,C). Given that cortactin employs its SH3 domain to engage Abl2 (Lapetina et al., 2009), as well as other synaptic proteins (Chen & Hsueh, 2012; MacGillavry et al., 2016; Mikhaylova et al., 2018), we mutated the conserved tryptophan residue in the cortactin SH3 domain to alanine (W525A), which disrupts its binding to the Abl2 Pro-X-X-Pro-X-X-Pro motif (Figure 3A). When introduced into cortactin KD neurons, the W525Acort^r^-RFP was depleted from spines as compared to WT cort^r^-RFP (Figure 3B,C). Thus, the cortactin SH3 domain is a major tether for its spine enrichment, while mutation of the three tyrosine residues does not impact cortactin enrichment in spines.

**Figure 3.**
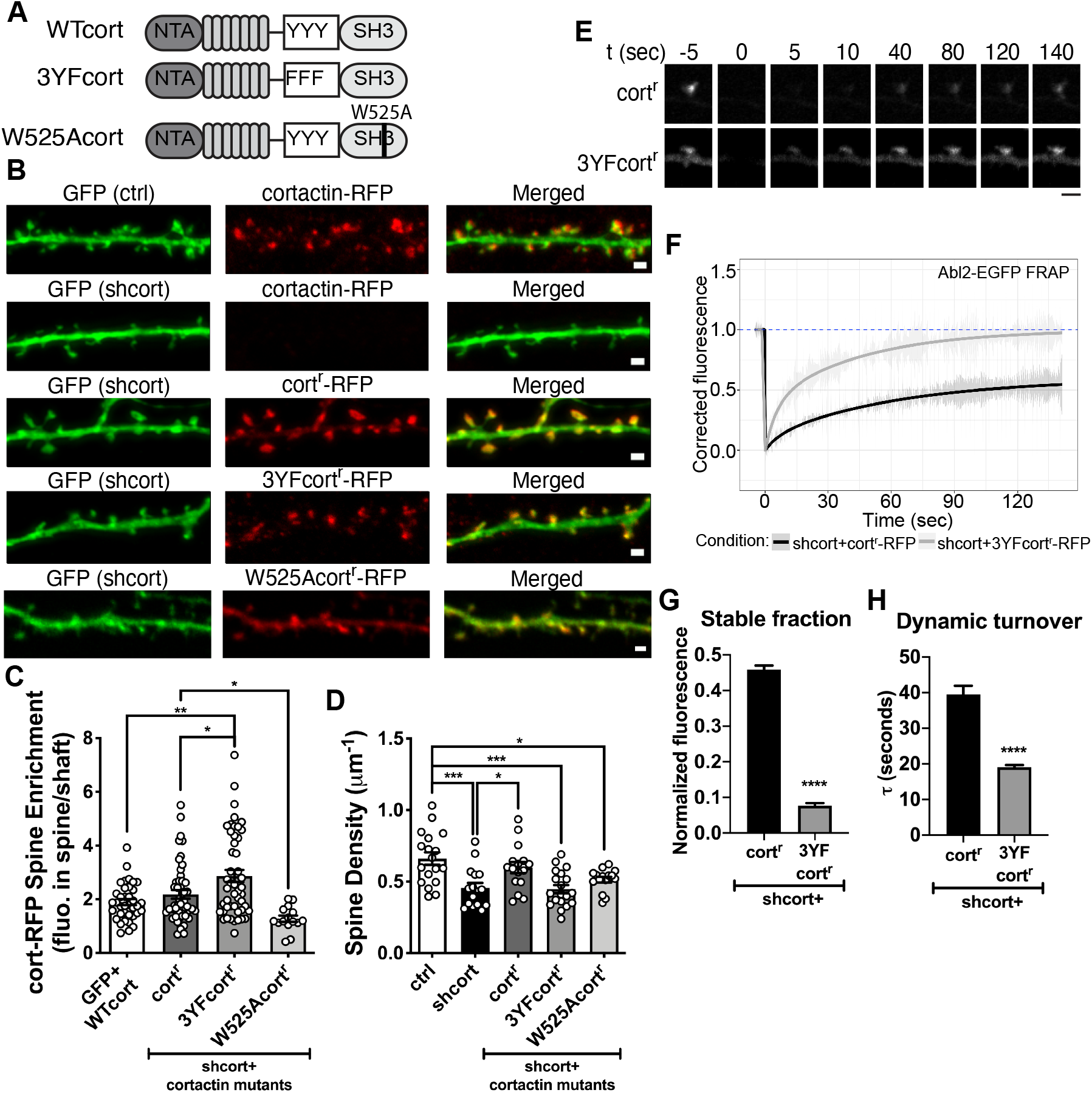
Cortactin phosphorylation is critical to support dendritic spine stability. **A.** Illustration of cortactin domain mutants used in the experiment. **B.** Representative confocal images of neurons transfected with GFP-tagged control vector or shcort. Cortactin KD neurons are co-transfected with cortactin-RFP, cort^r^-RFP, 3YFcort^r^-RFP or W525Acort^r^-RFP. Scale bar = 1 μm. **C.** The spine enrichment of cortactin-RFP, cort^r^-RFP, 3YFcort^r^-RFP or W525Acort^r^-RFP in control and cortactin KD neurons shows that cort^r^-RFP and 3YFcort^r^-RFP enrich in spines as well as cortactin-RFP. W525Acort^r^-RFP exhibits weak spine enrichment (mean = 1.278), but is significantly less than cort^r^-RFP. Data are means + SEM (n = 15-48 neurons/group). *p < 0.05, **p < 0.01 (ordinary one-way ANOVA with Tukey multiple comparisons test). **D.** Dendritic spine density of control neurons and cortactin KD neurons. Cortactin KD shows reduced spine density. Complementation with cort^r^-RFP, but not 3YFcort^r^-RFP or W525Acort^r^-RFP, restores spine density. Data are means + SEM (n = 15-19 neurons/group). *p < 0.05, ***p < 0.001 (ordinary one-way ANOVA with Tukey multiple comparisons test). **E.** Representative time-lapse confocal images of Abl2-EGFP co-expressed with cort^r^-RFP, or 3YFcort^r^-RFP in cortactin KD neurons. Only the Abl2-EGFP channel is shown. Time = 0 seconds indicates the frame immediately following photobleaching. Scale bar = 2 μm. **F.** Abl2-EGFP turnover was fit to a one-phase exponential curve. In the presence of 3YFcort^r^-RFP, Abl2 has a lower stable fraction **(G)** and faster turnover kinetics **(H)**. Data are means + SEM (n = 11 spines/group). ****p < 0.0001 (unpaired t test).

Cortactin KD neurons exhibited a 31 ± 8% reduction in spine density (Figure 3D), a slightly larger reduction to that previously reported (Hering & Sheng, 2003). Expression of W525Acort^r^-RFP, which did not enrich in spines, did not rescue this reduction in dendritic spine density (Figure 3D). Interestingly, even though it was *more* enriched in spines than cort^r^-RFP, 3YFcort^r^-RFP did not rescue the reduction in dendritic spine density resulting from cortactin KD (Figure 3D, summarized in Table 1). Phosphorylation of Y421 or Y466 is necessary for Abl2 binding to cortactin both *in vitro* and in cells, suggesting that this deficit in dendritic spine density could result from decreased Abl2:cortactin interaction in dendritic spines.

To test if disrupted cortactin phosphorylation impaired spine stability by impacting Abl2 tethering in spines, we co-expressed Abl2-EGFP with cort^r^-RFP or 3YFcort^r^-RFP in cortactin KD neurons and performed fluorescence recovery after photobleaching (FRAP) analysis on Abl2-EGFP (Figure 3E-H). When co-expressed with cort^r^-RFP a high proportion (45.9 ± 1.1%, Figure 3F,G) of Abl2-EGFP does not recover, indicating stable tethering in the spine. The dynamic Abl2-EGFP pool exhibits slow, one-phase recovery (τ = 39.5 ± 2.4 s, Figure 3F,H). However, co-expression of Abl2-EGFP with 3YFcort^r^-RFP resulted in a reduction of the stable pool of Abl2-EGFP (7.7 ± 0.7%, Figure 3F,G) and faster turnover of the dynamic proportion (τ = 19.0 ± 0.7 s, Figure 3F,H), indicating reduced Abl2 tethering within spines. These data, together with the findings that the kinase-inactive Abl2 and phospho-cortactin-binding defective Abl2-R198K mutant do not support normal spine density (Figure 1D) strongly suggest that Abl2-mediated cortactin phosphorylation and subsequent Abl2 binding to phospho-cortactin, promotes Abl2 tethering within the spine, thereby promoting dendritic spine stability (summarized in Table 1).

### Abl2 and cortactin are required to maintain the stable actin pool in dendritic spines

Cortactin-mediated enrichment of Abl2 into dendritic spines is critical to maintain normal dendritic spine density (Figure 1D and 3D). However, this raised the central mechanistic question of whether and how the regulation of actin polymerization and stability by Abl2 and/or cortactin is important to promote normal spine density. Thus, we asked whether alterations in these proteins impacted actin levels and turnover and how these alterations relate to changes in net dendritic spine density, as well as spine size. First, we measured filamentous actin levels using Atto647N-phalloidin staining of fixed cells and quantified regions of interest corresponding to dendritic spines, guided by expression of RFP that revealed neuronal structure (Figure 4A,B). Although filamentous actin levels were broadly distributed among spines, we did not observe shifts in the distribution of total actin levels under any condition (Figure 4B).

**Figure 4.**
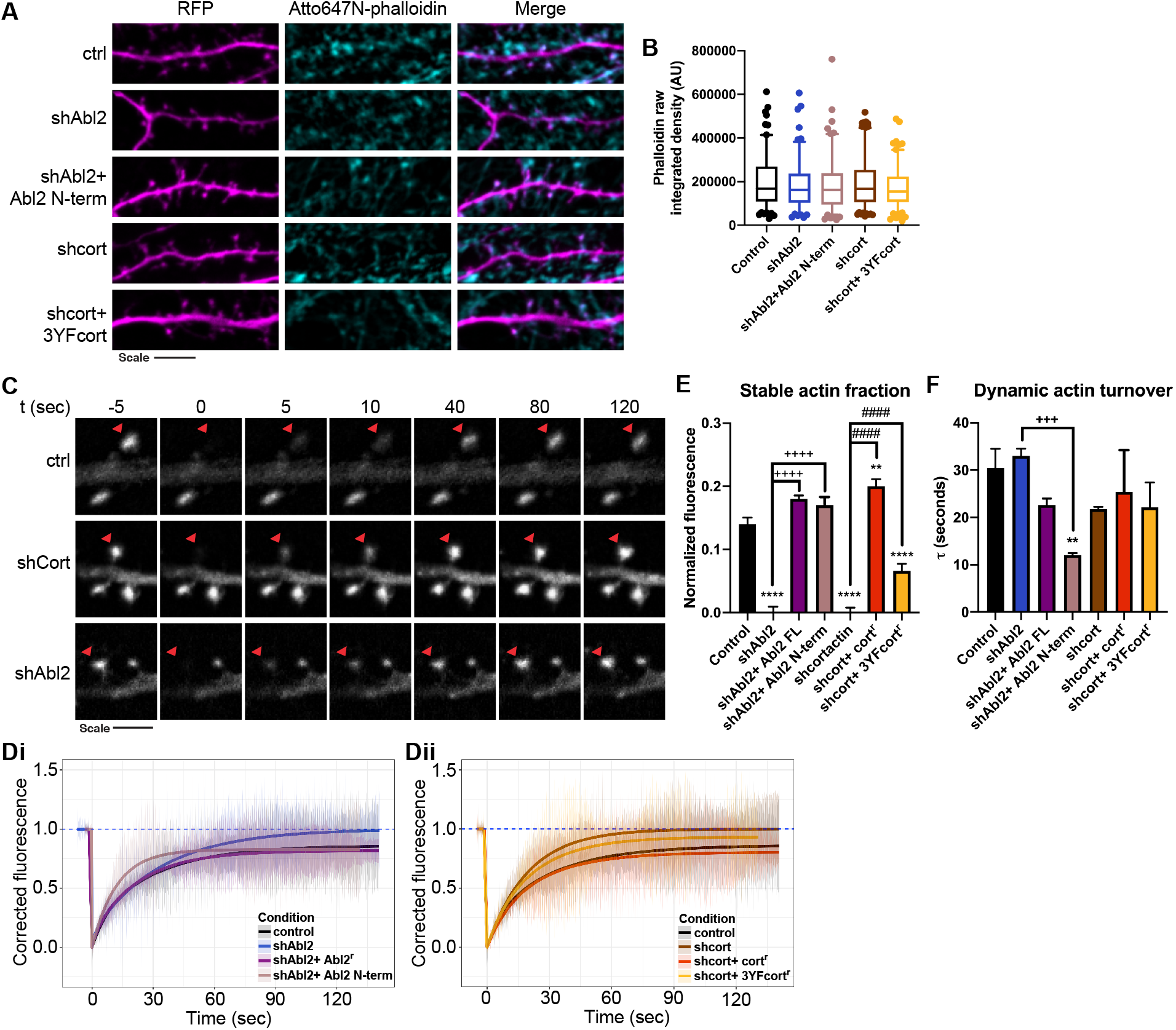
Abl2 and cortactin are required to maintain the stable actin pool in dendritic spines. **A.** Representative confocal images of neurons transfected with control vector, shAbl2, shAbl2 reconstituted with Abl2 mutants, shcort or shcort reconstituted with cortactin mutants. Neurons were stained for RFP and with a fluorescent phalloidin to visualize actin filaments in spines. Scale bar = 5 μm. **B.** Quantified intensity of fluorescent phalloidin. Data are displayed as box (25^th^ percentile, median and 75^th^ percentile) and whisker (5-95%) plots (n = 126-183 spines/condition). p > 0.999 (Kruskal-Wallis test with Dunn’s multiple comparison test). **C.** Representative time-lapse confocal images of GFP-actin expressed in dendritic spines of control neurons or neurons with Abl2 or cortactin KD. Time = 0 seconds indicates the frame immediately following photobleaching and the red arrowhead indicates the spine that was photobleached. Scale bar = 2 μm. **D.** GFP-actin FRAP was fit to a two-phase exponential recovery curve. Recovery curves are shown for neurons transfected with control vector, shAbl2 and shAbl2 reconstituted with Abl2^r^-RFP or Abl2 mutants **(Di)** or shcort and shcort reconstituted with cortactinr-RFP or cortactin mutants **(Dii)**. GFP-actin fluorescence does not recover fully in control neurons, but it reaches full recovery when Abl2 or cortactin is knocked down. The stable actin fraction **(E)** and dynamic actin turnover **(F)**, computed from the fitted curve is shown for each condition. Data are means + SEM (n = 16-28 spines/condition). **p < 0.01, +++p < 0.001, ****,++++,####p < 0.0001 (ordinary one-way ANOVA with Tukey multiple comparisons test).

We also performed FRAP analyses on GFP-actin to test whether manipulation of Abl2 or cortactin impacted the relative proportions of dynamic vs. stable actin content of the spine, and how this related to alterations in spine size or density (Figure 4C-F). Actin recovery time constants are best fit to a two phase recovery (Koskinen, Bertling, & Hotulainen, 2012; Star, Kwiatkowski, & Murthy, 2002): a fast component representing diffusion of GFP-actin monomers into the bleached spine (τ_monomer_ = ~1.5-3 s) and a slower component representing GFP-actin incorporation into the dynamic polymerizing actin network. The small unrecovered fraction represents stable actin that recovers on the order of 10s of minutes.

Consistent with previous reports (Koskinen et al., 2012; Koskinen & Hotulainen, 2014; Star et al., 2002), we found that 13.9 ± 1.0% of the GFP-actin fluorescence did not recover (stable fraction) after photobleaching (Figure 4C-E). Abl2 or cortactin KD results in a complete recovery of GFP-actin fluorescence (Figure 4D,E), indicating, at the population level, that loss of Abl2 and cortactin yield spines that lack a stable actin pool. Re-expression of Abl2^r^-RFP or cort^r^-RFP restored the pool of stable actin when expressed in Abl2 or cortactin KD neurons, respectively (Figure 4D,E). Interestingly, complementation of Abl2 KD neurons with Abl2 N-term-RFP, which rescued dendritic spine loss and cortactin enrichment in spines, also completely rescued the stable actin pool (Figure 4Di,E). Unexpectedly, rescue with the Abl2 N-term-RFP also increased the recovery rate of the dynamic pool (τ_control_ = 30.5 ± 4.1 s, τ_Abl2 N-term_ = 12.1 ± 0.4 s, Figure 4Di,F). This may result from the elevated expression levels and consequently elevated kinase activity of Abl2 N-term compared to full-length Abl2 (Peacock et al., 2007).

We next tested how disruption of cortactin phosphorylation impacts GFP-actin turnover. Complementation of cortactin KD neurons with 3YFcort^r^-RFP yielded partial rescue of the stable actin pool (Stable fraction = 13.9 ± 1.0% for control and 6.6 ± 1.1% for 3YFcort^r^-RFP, Figure 4Dii,E). 3YFcort^r^-RFP did not fully rescue dendritic spine density (Figure 3D), indicating that the remaining stable actin pool is not sufficient to stabilize spines. These data demonstrate that Abl2 and cortactin are novel key regulators of the stable actin pool in dendritic spines.

### Loss of Abl2 and cortactin differentially impact small versus large dendritic spines

Previous measurements of actin turnover in dendritic spines show that the amount of stable actin is proportional to the spine volume squared (Honkura et al., 2008). A subset of remaining spines following Abl2 depletion exhibited gradual spine head size enlargement (Y. C. Lin et al., 2013; Xiao, Levy, Rosenberg, Higley, & Koleske, 2016). These findings led us to examine how changes in actin pool sizes and recovery kinetics correlated with spine head size. For each spine analyzed, we measured the full-width at half-maximum (FWHM) of the spine head in the pre-bleach frames and took the average of those frames to approximate individual spine head widths (Figure 5A). The distribution of spine head widths of photobleached spines was significantly broader in the Abl2 KD condition than in control spines (Figure 5B), consistent with measurements made previously (Y. C. Lin et al., 2013). In notable contrast, the distribution of spine head widths for cortactin KD photobleached spines were not significantly different from control spines (Figure 5C). These differences in the populations of spine head widths raise the question of whether spines of different head sizes exhibit differences in the stable pool in Abl2- or cortactin- KD neurons.

**Figure 5.**
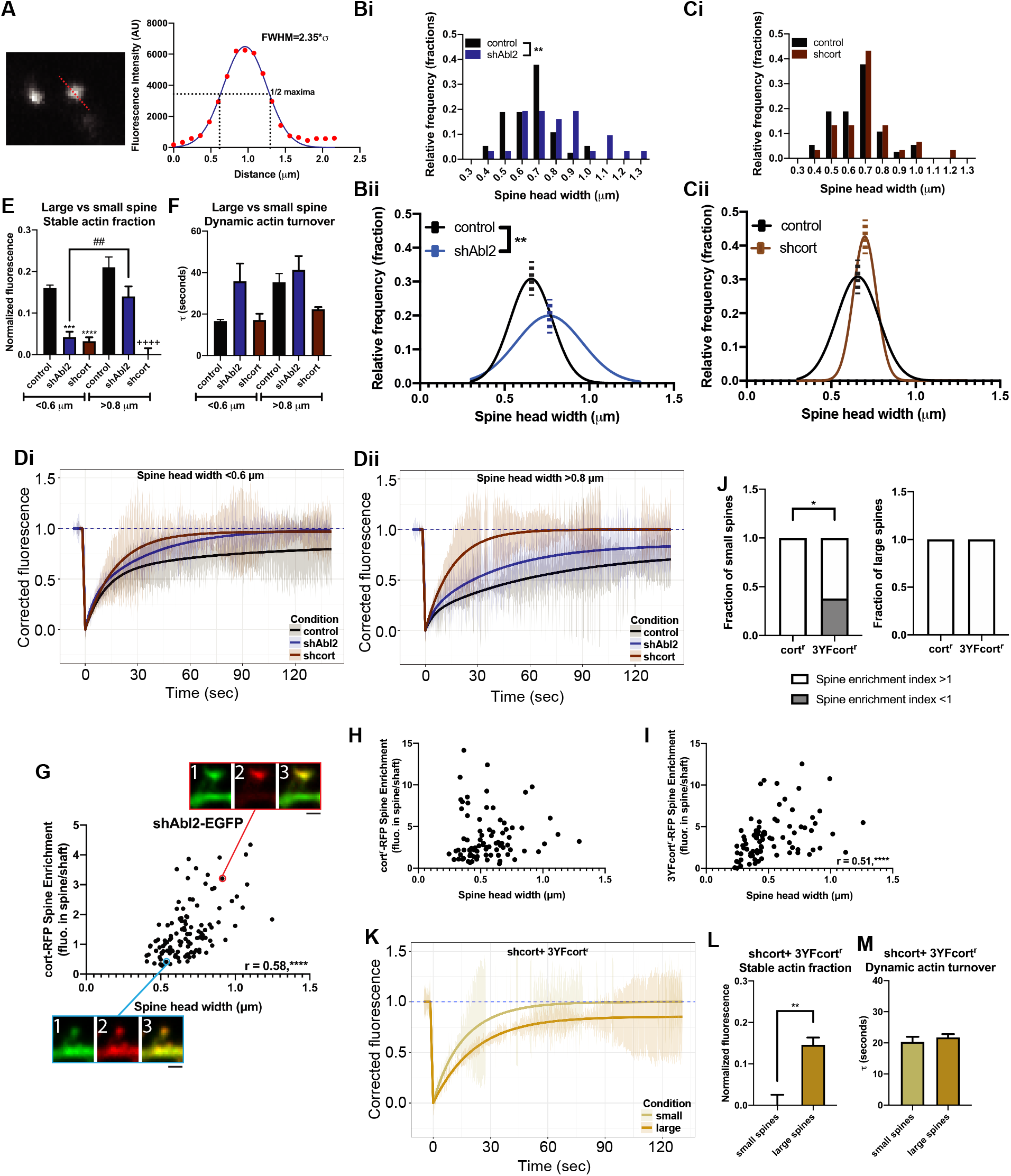
Loss of Abl2 and cortactin differentially impact small versus large dendritic spines. **A.** FWHM was measured for each spine head subjected to photobleaching by plotting the fluorescence profile across a line and fitting to a Gaussian curve. **B.** The distribution of the average spine head width of each photobleached spine is shown both as a frequency distribution **(Bi, Ci)** and fit to a Gaussian curve **(Bii, Cii)**. Center of the curves are denoted with a dotted line. The shAbl2 spine heads subjected to photobleaching are significantly enlarged compared to control spines (control, n = 26; shAbl2, n = 24 spines). **p < 0.01 (welch’s t test). To account for any differences in populations of spine heads that were photobleached, data was parsed into small spines < 0.6 μm **(Di)** and large spines > 0.8 μm **(Dii)** for control, shAbl2 and shcort data and recovery curves were generated. Difference in the stable actin fraction **(E)** and dynamic actin turnover **(F)** is shown. Data are means + SEM (n = 6-13 spines/condition). ##p < 0.01, ***p < 0.001, ****,++++p < 0.0001 (ordinary one-way ANOVA with Tukey multiple comparisons test). **G.** Scatterplot for spine head width versus cort-RFP spine enrichment in Abl2 KD neurons shows a positive correlation (n = 117 spines). r = 0.58, ****p < 0.0001 (Pearson correlation test). Representative confocal images are shown for a small spine (bottom), demarked with a blue outline, and a large spine (top), demarked with a red outline. Channels: (1) shAbl2-EGFP, (2) cort-RFP and (3) merged. Scale bar = 1 μm. **H.** Scatterplot for spine head width and cort^r^-RFP spine enrichment in cortactin KD neurons shows no correlation (n = 83 spines). r = 0.19, p = 0.078 (Pearson correlation test) **I.** Scatterplot for spine head width and 3YFcort^r^-RFP spine enrichment in cortactin KD neurons shows a positive correlation (n = 83 spines). r = 0.51, ****p < 0.0001 (Pearson correlation test). **J.** The fraction of small and large spines that show preferential spine enrichment (spine enrichment > 1) from the population of cort^r^-RFP and 3YFcort^r^-RFP spines. A significant fraction of small 3YFcort^r^-RFP spines do not have a preferential enrichment in spines (spine enrichment < 1) compared to small cort^r^-RFP spines (small spines: n = 10 cort^r^-RFP spines, 21 3YFcort^r^-RFP spines; large spines: n = 11 cort^r^-RFP spines, 14 3YFcort^r^-RFP spines). *p < 0.05 (Fisher’s exact test). **K.** Due to a significant correlation between spine head width and 3YFcort^r^-RFP spine enrichment, and differences in enrichment between small and large spines, data for GFP-actin FRAP in shcort+ 3YFcort^r^-RFP neurons was parsed into small spines < 0.6 μm and large spines > 0.8 μm and recovery curves were generated. Difference in the stable actin fraction **(L)** are shown with no changes in the dynamic actin turnover **(M)**. Data are means + SEM (n = 5 spines/condition). **p < 0.01 (t test).

To test how the relative size of the stable actin pool related to spine size, we segmented our FRAP dataset based on spine head widths and classified small spines based on FWHM < 0.6 μm and large spines with FWHM > 0.8 μm, less than and greater than one standard deviation from the control mean, respectively. Small and large spines in the control and cortactin KD datasets represent approximately 25% of the total population of spines analyzed in Figure 4. However, for the Abl2 KD dataset, small spines account for approximately 20% of the total population, while large spines are 40%. Small Abl2 KD spines, as well as both small and large cortactin KD spines, exhibited significantly smaller stable actin pools as compared to their respective control (Figure 5D,E), consistent with the pooled data in Figure 4D,E. Dynamic actin turnover rates were not significantly changed in any of these conditions (Figure 5F). The stable pool size trended toward an increase in larger spine head widths, consistent with a previous report (Honkura et al., 2008) (Figure 5Dii,E), although we could not establish significance in our data set due to the small number of large spines in the control neuron comparison group. Large Abl2 KD spines had a significantly larger stable actin pool than small Abl2 KD spines, and this was not significantly different from the stable actin pool size in large control spines (Figure 5Dii,E). Together, these data suggest that while cortactin is absolutely required to maintain the stable actin pool in all spines, the subset of large spines in Abl2-deficient neurons retain the stable actin pool.

The remaining stable actin pool in large Abl2 KD spines suggested that there may be an Abl2-independent enrichment of cortactin in these large spines, that enables actin stabilization. To determine if cortactin enrichment in Abl2 KD spines correlates with spine head size, we quantified the FWHM and cort-RFP spine enrichment for spines on Abl2 KD neurons (Figure 5G). We found a positive Pearson Correlation (r = 0.58) between spine size and cortactin enrichment. In cortactin KD neurons reconstituted with cort^r^-RFP, there is no significant correlation between spine head width and cortactin spine enrichment (Figure 5H). This suggests that elevated cortactin levels in large Abl2 KD spines are sufficient to support a stable actin pool and protect these spines from destabilization.

The 3YFcort mutant is enriched in spines to a greater extent than WT cort (Figure 3C), but it untethers Abl2 from spines (Figure 3F-H) and only partially rescues the stable actin pool (Figure 4Dii,E), resulting in a lower spine density (Figure 3D). Therefore, we used the parsing method above to document and quantify the Abl2-independent enrichment of cortactin, using 3YFcort^r^-RFP, and how this correlated with actin stability in large versus small spines of Abl2 KD neurons. Similar to cortactin spine enrichment in Abl2 KD neurons, 3YFcort^r^-RFP spine enrichment is positively correlated with spine head size (Pearson correlation r = 0.51) (Figure 5I). Furthermore, a significant proportion of small spines do not show spine enrichment of 3YFcort^r^-RFP (Figure 5J). Given differential enrichment of 3YFcort in small versus large spines, we performed GFP-actin FRAP in these small and large spines (Figure 5K). While in aggregate, the 3YFcort^r^-RFP data had a partial rescue of the stable actin pool (Figure 4Dii,E), we found that small spines had no stable actin pool, similar to cort KD neurons, and large spines had stable actin pools comparable in size to spines in control neurons (Stable fraction = 14.6 ± 1.8%, Figure 5L). No significant changes in the dynamic actin turnover were observed in the cort KD + 3YFcort^r^-RFP spines irrespective of size (Figure 5M). These data indicate that even while cortactin tethering of Abl2 is important for maintaining small dendritic spines, in the absence of Abl2, cortactin associates with another determinant in the spine that scales with spine size.

### Tonic enhancement of synaptic activity rescues cortactin depletion and spine loss in Abl2 KD neurons

Synaptic activity bidirectionally reorganizes the molecular composition and structure of dendritic spines. Actin is a major substrate for bidirectional plasticity, as changes in actin dynamics rapidly correlate with changes in dendritic spine structure (Fukazawa et al., 2003; Okamoto, Nagai, Miyawaki, & Hayashi, 2004). Moreover, manipulations of activity modulate the translocation of cortactin and other synaptic proteins into and out of the spine (Bosch et al., 2014; Hering & Sheng, 2003; Iki et al., 2005; K. Kim et al., 2015). Given the impact of Abl2 and cortactin on the stable actin pool (Figure 4, 5) and this unique link between spine head size and cortactin spine enrichment in Abl2-deficient neurons (Figure 5G-M), we asked whether tonic changes in activity impacted these phenotypes.

During the duration of Abl2 KD, we treated cultured neurons with either 1 μM tetrodotoxin (TTX) or 20 μM bicuculline to ask how chronic blockade or enhancement of activity, respectively, modulated dendritic spine density, cortactin localization and gross changes in dendritic spine structure (Figure 6A). There was no significant difference in the dendritic spine density of TTX and bicuculline treated control neurons (Figure 6C). TTX did not impact Abl2 deficient neurons – compared to control transfected cells, they had a significantly reduced dendritic spine density (Figure 6B,C; left) and a 31 ± 10% reduced spine enrichment of cortactin (Figure 6B,D; left), similar to untreated Abl2-deficient neurons (Figure 2E). Interestingly, tonically increasing activity with bicuculline rescued the reduced spine density in Abl2 KD neurons (Figure 6B,C; right) and maintained cortactin enrichment in these spines (Figure 6B,D; right). Bicuculline treatment also normalized the spine head width distribution in Abl2 KD neurons to mirror the head widths observed in control neurons – the distribution of spine head widths was no longer right shifted (Figure 6E, right), reflecting that loss of Abl2 no longer results in a significant enlargement of spine heads. These data indicate that tonically increasing activity compensates for Abl2 deficiency. TTX did not impact net cortactin depletion from spines or net spine loss in Abl2 KD neurons, but it normalized spine head size (Figure 6E, left). Consistent with normalization of dendritic spine head widths in Abl2 KD neurons by both TTX and bicuculline, we no longer observed the correlation between spine head width and cortactin spine enrichment (Figure 6F) observed in Abl2 KD neurons in basal conditions (Figure 5G).

**Figure 6.**
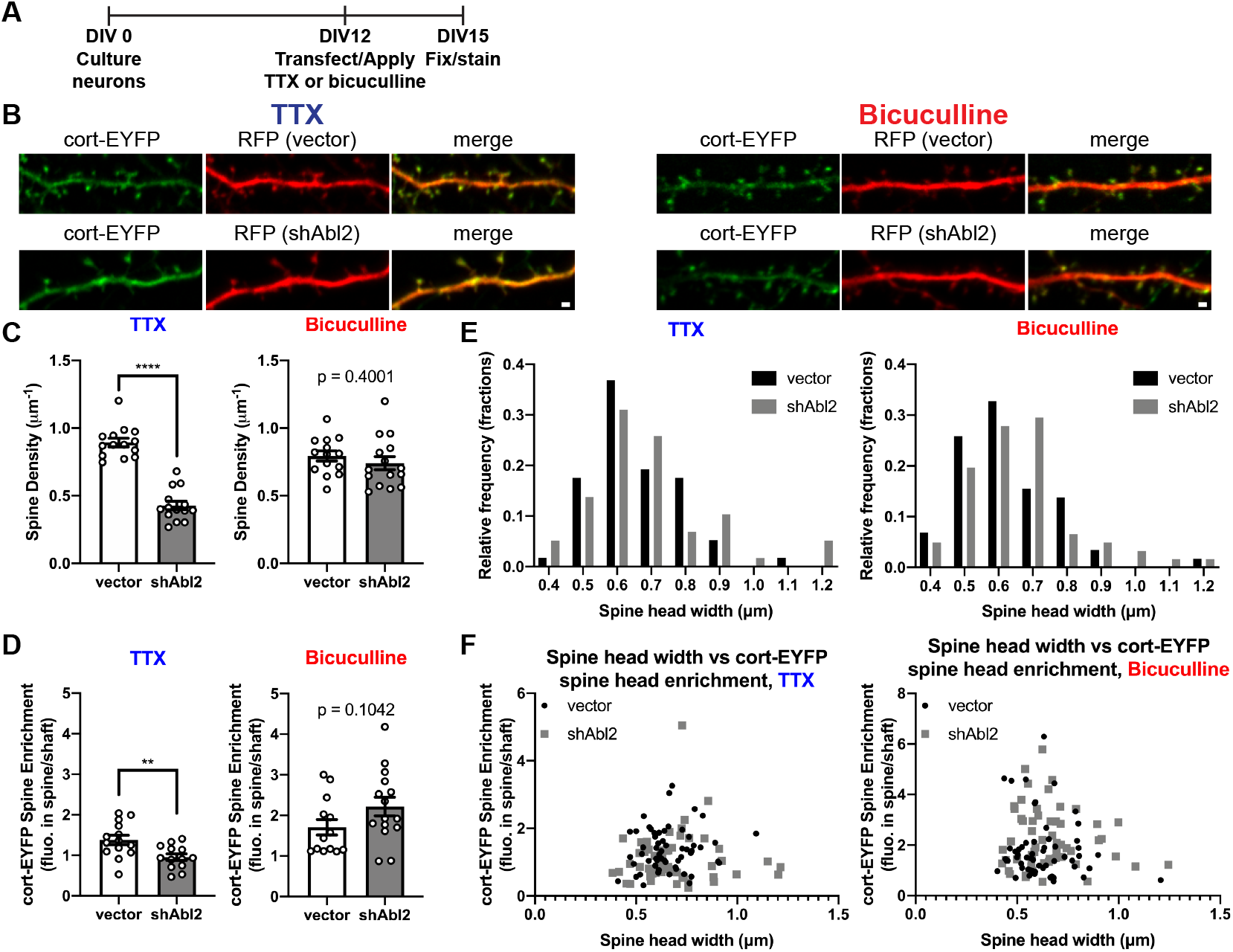
Tonic enhancement of synaptic activity rescues cortactin depletion and spine loss in Abl2 KD neurons. **A.** Experimental set up. Neurons were cultured for 12 days before knocking down Abl2. During the course of knockdown, cells were treated with 1 μM tetrodotoxin (TTX) or 20 μM bicuculline methiodide for 72 hours before fixation and staining for YFP and RFP, to visualize cortactin localization and cell morphology, respectively. **B.** Representative confocal images of neurons co-transfected with control vector or shAbl2 and cortactin-EYFP. Scale bar = 1 μm. **C.** Dendritic spine density of control and Abl2 KD neurons treated with TTX or bicuculline. TTX treated neurons show a reduced dendritic spine density upon Abl2 KD, similar to basal conditions. However, treatment of neurons with bicuculline rescues dendritic spine deficits upon Abl2 KD. Data are means + SEM (n = 14-15 neurons/condition). ****p < 0.0001 (unpaired t test). **D.** Spine enrichment of cortactin-EYFP in control and Abl2 KD neurons treated with TTX and bicuculline. Note that cortactin enrichment in spines is retained in Abl2 KD neurons treated with bicuculline. Data are means + SEM (n = 13-15 neurons/condition). **p < 0.01 (unpaired t test). **E.** The distribution of spine head widths in control and Abl2 KD conditions treated with TTX and bicuculline is shown as a frequency distribution. Abl2 KD spines do not show an enlargement compared to control spines when treated with TTX or bicuculline (n = 57-61 spines/condition). TTX, vector vs shAbl2 p = 0.7434; bicuculline, vector vs shAbl2 p = 0.2269 (Mann-Whitney test). **F.** Scatterplot for spine head width and cort-EYFP spine enrichment in control and Abl2 KD neurons treated with TTX and bicuculline does not show a positive correlation for any of the conditions (n = 57-61 spines/condition).

## Discussion

Reductions in dendritic spine density are associated with cognitive, perceptual, and affective deficits in psychiatric, neurodevelopmental, and neurodegenerative disorders. The actin cytoskeleton, consisting of both dynamic and stable pools, is a major determinant of spine structure and stability. Here, we identify Abl2 and cortactin as key regulators of the stable actin pool in dendritic spines. We show that disruption of Abl2 SH2 domain binding to phosphorylated cortactin reduces enrichment of both proteins in spines and disrupts the stable actin pool. Loss of the stable actin pool is associated with significant reductions in spine density. Curiously, we also find that a small subset of larger spines in Abl2-deficient neurons exhibit increased cortactin recruitment and a stable actin pool, suggesting that cortactin acts via both Abl2-dependent and independent mechanisms. Together our results define new key regulators of the stable actin pool in spines and demonstrate the importance of stable actin in regulating spine shape and stability.

### A role for stable actin in dendritic spine stability

Consistent with previous measurements, we find that only a small fraction of spine actin is stable (Fukazawa et al., 2003; Honkura et al., 2008; Koskinen et al., 2014; Star et al., 2002). This stable actin pool appears to localize to the spine base (Fukazawa et al., 2003), where cortactin has also been shown to concentrate (Racz & Weinberg, 2004). We demonstrate that depletion of the stable actin pool is tightly associated with decreased dendritic spine stability and overall decreased spine density. Although stable actin is not very abundant in dendritic spines, it seems to play a major role in dendritic spine stability, raising the fundamental question of how it contributes to spine homeostasis.

The ongoing maintenance of spine shape and synaptic transmission requires continual dynamic actin polymerization and turnover (Honkura et al., 2008; C. H. Kim & Lisman, 1999; K. Kim et al., 2015; Krucker, Siggins, & Halpain, 2000). It is possible that these dynamic actin filaments use the stable actin pool as a base for their nucleation. In maturing synapses, this dynamic pool also undergoes continual retrograde flow (Koskinen et al., 2014), which likely exerts force on the stable pool at the spine base. Here, the stable pool may play a supporting role to resist this force and stabilize the spine. The size of the stable pool scales with spine size (Honkura et al., 2008; Koskinen et al., 2012), consistent with possible roles as a support structure for the peripheral dynamic pool. Additionally, large bundles of actin filaments are enmeshed with the spine apparatus in a subset of spines (Capani, Martone, Deerinck, & Ellisman, 2001). Although the function of the spine apparatus remains enigmatic, dendritic spines containing a spine apparatus are larger and have more potent synapses (Holbro, Grunditz, & Oertner, 2009).

Furthermore, only spines that contain a spine apparatus grow during cLTP (Borczyk, Sliwinska, Caly, Bernas, & Radwanska, 2019; Deller et al., 2003). Therefore, the stable actin pool may support the spine apparatus to promote spine stability. Previous work from Mikhaylova et al. demonstrated caldendrin depletion also results in a loss of the stable actin pool and spine plasticity deficits (Mikhaylova et al., 2018). Furthermore, caldendrin depletion enhances cortactin turnover in dendritic spines, suggesting that these changes in spine actin stability may be the indirect result of cortactin loss from spines.

### How do Abl2 and cortactin maintain the stable pool of actin in spines?

We show here that Abl2 and cortactin are essential to maintain the stable actin pool in dendritic spines. Although Abl2 and cortactin can each independently bind and stabilize actin filaments, they synergize at much lower binding stoichiometries *in vitro* (Courtemanche et al., 2015; Scherer et al., 2018). The Abl2 N-terminal half, which neither binds actin nor promotes cortactin binding to actin, is sufficient to support the stable actin pool in Abl2-deficient neurons. The Abl2 kinase domain, located in the protein’s N-terminus, phosphorylates cortactin, creating a binding site for the Abl2 SH2 domain. Mutation of this interface, either via point mutations to the Abl2 SH2 domain or the cortactin phosphorylation sites, reduces the stable actin pool and leads to spine loss. We hypothesize that Abl2 phosphorylation and tethering to cortactin keeps it in the spine, promoting interactions with actin and possibly other proteins that are critical for it to maintain the stable actin pool. We also find that the nonphosphorylatable (3YF) cortactin is more enriched in spines, and Abl2 is more dynamic in spines reconstituted with 3YFcort. Uncoupling from Abl2 may free up 3YFcortactin to interact with other binding partners in the spine, including, but not limited to, the Shank family of proteins (MacGillavry et al., 2016).

We also found that rescue of Abl2 with the Abl2 N-terminus yields much faster dynamic actin recovery. The Abl2 N-terminus expresses at elevated levels over Abl2, yielding increased Abl2 kinase activity (Peacock, Couch, & Koleske, 2010). In addition to cortactin, Abl family kinases phosphorylate N-WASp and WAVE2 (M. M. Miller et al., 2010; Stuart, Gonzalez, Kawai, & Yuan, 2006), which can both activate the Arp2/3 complex (Martinez-Quiles, Ho, Kirschner, Ramesh, & Geha, 2004). Abl2 also phosphorylates the RhoA inhibitor p190RhoGAP (Sfakianos et al., 2007), which attenuates RhoA/ROCK signaling and could activate actin severing by cofilin (Bravo-Cordero et al., 2011). Increasing Arp2/3-mediated actin branching and cofilin severing would both result in increases in dynamic turnover.

### How do large Abl2-deficient spines retain a stable actin pool?

Depletion of Abl2 results in a significant loss of cortactin from spines and overall spine loss (Y. C. Lin et al., 2013; Xiao et al., 2016). Curiously, while net spine stability is decreased in Abl2-deficient neurons, a subset of the remaining spines enlarge (Figure 5). These large spines retain a stable actin pool and are enriched for cortactin. Previous work has also shown that the enlarged subset of spines in Abl2-deficient mice have larger GluN2B-mediated NMDAR currents (Xiao et al., 2016). We find that tonically increasing synaptic activity with bicuculline rescues cortactin spine enrichment in Abl2 KD neurons and overall dendritic spine stability (Figure 7) and blocking presynaptic release with TTX prevents spine head enlargement. Paradoxically, other studies have shown NMDAR stimulation causes cortactin to redistribute from spines to dendrite shafts (Hering & Sheng, 2003; Iki et al., 2005), although it is unclear which class of NMDA receptors trigger cortactin exit from spines. It is clear that in the absence of Abl2, some large spines retain the ability to maintain cortactin levels and a stable actin pool, suggesting that cortactin may respond differently to NMDARs, possibly dependent on their subunit composition. In addition to Abl2, cortactin interacts via its SH3 domain with the synaptic scaffolds SHANK2 (MacGillavry et al., 2016) and cortactin-binding protein 2 (CTTNBP2) (Chen & Hsueh, 2012), suggesting that these large spines may tether cortactin by either of these proteins. Also, the cortactin repeats region of cortactin, which binds and stabilizes actin filaments (Scherer et al., 2018; Weed et al., 2000), is sufficient to localize cortactin in spines (Hering & Sheng, 2003), suggesting that interactions with actin may be sufficient. Neurotrophic cues, such as BNDF, have been proposed to drive cortactin into spines (Iki et al., 2005), where it would be able to interact with actin filaments. Therefore, retaining cortactin localization in spines via tethering to various synaptic scaffolds or even actin itself likely compensates for a loss of Abl2 and its disruption of the stable actin pool.

**Figure 7.**
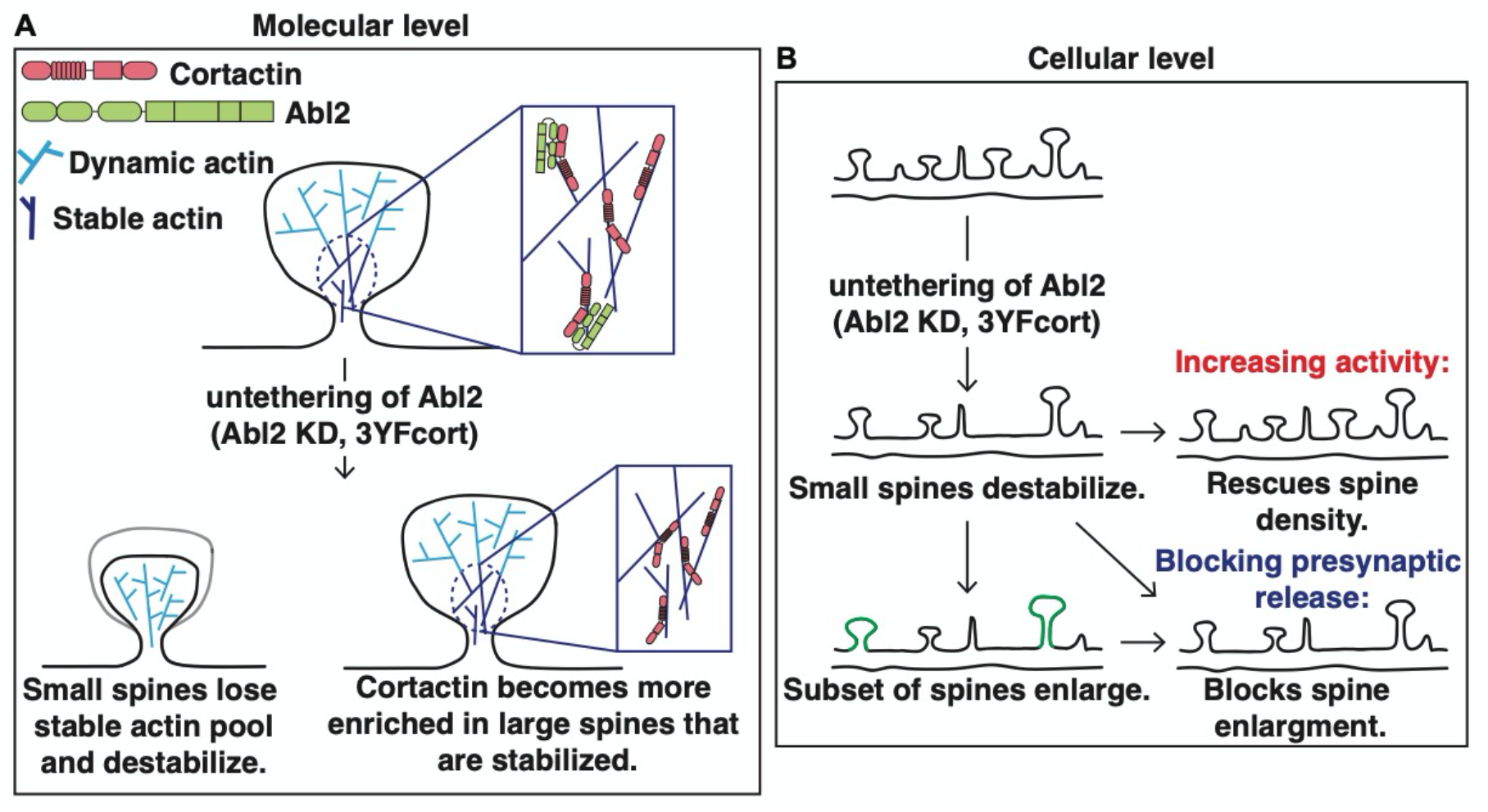
Model of Abl2:cortactin regulation of dendritic spine stability. **A.** At the molecular level, a subset of dendritic spine actin is stable (dark blue), while the majority is highly dynamic (light blue). The Abl2 SH2:phospho-cortactin interaction is essential in maintaining this pool of stable actin (top, inset). Dendritic spine loss upon Abl2 KD is accompanied by loss of the stable actin pool in these small spines (bottom, left) and the redistribution of cortactin to large spines that retain a significant stable actin pool (bottom, right), by an Abl2-independent mechanism. **B.** A cellular perspective of how Abl2 and cortactin impact dendritic spines. At homeostasis, dendritic spines are all different sizes and maintain different neurotransmitter and synaptic machinery. Loss of Abl2 results in the gradual loss of cortactin from small spines and their eventual destabilization. Spine loss can be prevented in Abl2 KD neurons by tonically increasing activity. Furthermore, spine loss is accompanied by the enlargement of some remaining spines, which can be prevented by blocking presynaptic release.

### A model for Abl2 and cortactin in regulation of spine stable actin

While dendritic spines contain two kinetically distinct pools of spine actin, the function of the stable actin pool has been largely understudied. Our data indicate that phosphorylated cortactin tethers Abl2, via its SH2 domain, in spines and this interaction is essential in maintaining the stable actin pool (Figure 7A). Upon loss of Abl2 or mutational disruption of phospho-tyrosine residues on cortactin, cortactin is lost from small dendritic spines and becomes enriched in a subset of large spines that retain the stable actin pool. Together, our data strongly support a model in which Abl2:cortactin interactions are critical for maintenance of the spine stable actin pool.

Our data also reveal interesting crosstalk between synaptic activity and the regulation of spine stability via the stable actin pool. Tonic enhancement of synaptic activity in Abl2-deficient neurons protects against cortactin depletion from spines and net spine loss. This observation points to a key activity-dependent mechanism that promotes cortactin enrichment in spines to stabilize them (Figure 7B). Gradual spine loss over days in Abl2-deficient neurons is accompanied by an enlargement of a subset of remaining spines (this study; (Y. C. Lin et al., 2013)). Disrupting activity, with TTX, in Abl2 KD neurons does not protect against spine loss, but it blocks enlargement, again pointing to an activity-driven process for enriching cortactin in spines. We note that the residual spines that remain on cortactin KD neurons do not enlarge, suggesting a direct role for cortactin in spine enlargement. These observations identify the stable actin pool as a key target of activity-mediated control of net spine size and stability.

## Materials and Methods

### Animals

All animal procedures were compliant with federal regulations and approved by the Animal Care and Use Committees at Yale University, University of Maryland at Baltimore and Hussman Institute for Autism. Balb/c mice were purchased from Charles River Laboratories. Animals were housed and cared for in a Yale-sponsored OPRR approved animal facility. Neonatal mice of both sexes were sacrificed for neuronal culture preparation.

### Cell culture and transfections

HEK293 cells were plated in 6-well plates and maintained in high glucose DMEM (Invitrogen) growth media supplemented with 1% pen/strep (Gibco), 2 mM L-glutamine (Gibco), and 10% fetal bovine serum (Sigma-Aldrich). Primary neuronal cultures were prepared from postnatal day 1 mouse hippocampus as previously described (Y. C. Lin et al., 2013). Neurons were maintained in serum free media containing 1% pen/strep, 2 mM L-glutamine, and 2% B27-supplement (Gibco) in Neurobasal media (NB-SFM) (Invitrogen). Glass coverslips in 24-well plates were coated with 20 μg/ml of poly-(D)-lysine (Corning Inc.) overnight at 37°C and 1 μg/ml laminin 111 (Corning Inc.) for 2 hours at 37°C. Neurons were plated at a cell density of 0.3×10^6^ cell/well. Transfection was performed using a calcium-phosphate precipitation method in both HEK293 cells and primary neurons. Tetrodotoxin citrate (1 μM; Tocris) and 20 μM bicuculine methiodide (Sigma-Aldrich) were added directly to the culture media immediately following transfection and incubated with neurons 72 h before fixation.

### Plasmids

Wild type and Abl2 mutants were constructed in N1-mRFP plasmid as previously described (Lapetina et al., 2009; A. L. Miller, Wang, Mooseker, & Koleske, 2004; Peacock et al., 2010). The shRNA-resistant mutations (Y. C. Lin et al., 2013) were introduced in WT-Abl2, Abl2-KI, Abl2 N-term, and Abl2-R198K mutants and subcloned into shAbl2 plasmid for the experiment shown in Figure 1. The shRNA targeted to mouse cortactin was generated by synthesizing two complementary oligonucleotides (shcort#1: 5’-tgcactgctcacaagtggacttcaagagagtccacttgtgagcagtgcttttttc-3’ and shcort#2: 5’-tcgagaaaaaagcactgctcacaagtggactctcttgaagtccacttgtgagcagtgca-3’) and cloned into the pLL3.7 vector (Rubinson et al., 2003) between HpaI and XhoI sites. Three tyrosine sites at 421, 466, and 482 were mutated to phenylalanine to make 3YFcortactin mutant as described previously (Lapetina et al., 2009). The shRNA resistant cortactin cort^r^-RFP, 3YFcort^r^-RFP and W525Acort^r^-RFP constructs were made by mutating three nucleotides in the shCort recognition sequences with two PCR primers: 5’-cgaagctttccaagcactgctctcaagtcgactccgtccggg-3’ and 5’-ccgaagccccggacggagtcgacttgagagcagtgcttgg-3’.

### Western Blot Analysis

Cells were harvested then lysed in buffer containing 20 mM Tris (pH7.5), 150 mM NaCl, 2 mM EDTA, 1% Triton X-100, and phosphatase and protease inhibitor cocktails. Protein concentration was determined using a Pierce BCA Protein assay Kit (Thermo Scientific). 75 μg of total protein from HEK293 cell lysates were used in immunoblots to determine the knockdown efficiency. Protein samples were run on a 10% SDS-PAGE and transferred to nitrocellulose membranes for immunoblotting with antibodies against RFP (Invitrogen) and β-actin (Abcam) in TBST containing 3% milk. Densitometry of blotted protein was performed on scanned films using ImageJ (NIH).

### Immunocytochemistry and morphometric analysis

Neurons were fixed with 4% paraformaldehyde in PBS and permeabilized with 0.1% Triton X-100 and 3% BSA in PBS. Cells were immunostained with primary antibodies against GFP (Rockland) or RFP (Chemicon) followed by Alexa488 and 594 conjugated secondary antibodies (Molecular Probes). For total filamentous actin staining with fluorescent-phalloidin, neurons were fixed with 2% PFA in cytoskeleton buffer (10 mM MES pH 6.8, 138 mM KCl, 3 mM MgCl_2_, 2 mM EGTA and 0.32 mM sucrose). Fixed cells were permeabilized with 0.3% TX100/TBS (150 mM NaCl, 20 mM Tris pH 7.4) and blocked with 0.1% TX100/2% BSA/10% Normal Donkey Serum/TBS. Cells incubated at 4° C overnight with anti-RFP (Rockland). The cells were then sequentially incubated with Alex594 conjugated secondary antibodies (Molecular Probes) and Atto647N-phalloidin (Sigma). For dendritic spine analysis, 2-3 dendrite segments per neuron were selected randomly from secondary or tertiary branches on the apical dendrite for imaging. Images were taken using a LSM710 confocal microscope with 63x NA 1.4 objective and 4x zoom. Dendritic spine density was quantified using ImageJ (NIH). Quantified dendrite segments were averaged for each neuron analyzed and each averaged dendritic spine density for a cell is represented as a sample (n) in the graphs. To measure the spine enrichment of cortactin and Abl2 content, a region of interest (ROI) was selected in the dendritic spine, adjacent dendrite shaft, and over a region in the background. Spine enrichment was quantified using the following equation:

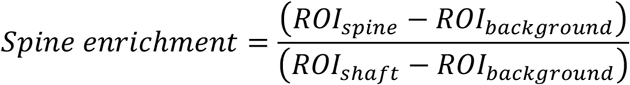

Spine enrichment was quantified for 4-6 spines per neuron then an average was taken for the cell. The average spine enrichment for a neuron is represented as a sample (n) in the graphs.

### Fluorescence Recovery After Photobleaching

Dissociated P0-1 mouse hippocampal neurons were plated onto 35-mm glass bottom dishes (MatTek). Cells were transfected using a calcium-phosphate precipitation method at 12 DIV with a 1:1 ratio of pN1-EGFP-human beta-Actin to pLL3.7-shRNA-mRFP plasmids targeting either Abl2 or cortactin. Live-cell FRAP imaging of pyramidal neurons in NB-SFM without phenol red culture medium was performed at 15 DIV – 72 hours after transfection, when we see net spine destabilization in Abl2 KD neurons (Y. C. Lin et al., 2013) – at 37°C using a 60x oil immersion objective on an UltraVIEW VoX (Perkin Elmer) spinning disc confocal microscope equipped with the PhotoKinesis (PK) Device. Volocity software (Improvision, Coventry, UK) was used to select the bleach ROI over the head of a spine on the secondary or tertiary branch of the apical dendrite. Images were acquired at 3 frames per second (fps) for 6-10 frames prior to bleaching. Prebleach fluorescence intensities were averaged and normalized to 1, and the curves were fit with two-phase exponential equations, as described with some modifications detailed below (Koskinen et al., 2014; Star et al., 2002). Prior to bleaching, 6-10 frames were captured then bleaching was performed using the 488 nm laser at 20% power with the following Ultraview PK settings: 100 PK cycles, 200 ms spot period and 1 PK spot cycle. During the recovery phase images were acquired at 3 fps for 10 seconds then for the duration of the experiment at 1 fps to increase sample protection and reduce phototoxicity on the sample. 3-5 spines/cell were bleached. Background fluorescence was determined in an area outside the cell using a ROI size equal to the bleach ROI. Potential FRAP-unrelated bleaching effects were corrected using a ROI with the same dimensions as the bleach ROI placed in an unbleached area with similar initial fluorescence as the bleach ROI. Fluorescence recovery (mobile fraction) over time was calculated by the following two-phase exponential equation:

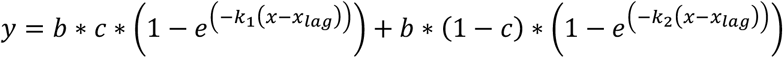

where *b* is the plateau, *c* is the fraction of dynamic actin, *k*_1_ (and *k*_2_) are rate constants with the units inverse seconds and *x_lag_*, is the time immediately following bleaching. The stable actin fraction is then defined as 1-*b* and τ, the time constant for the dynamic actin turnover, is defined as 1/*k*_1_ and has units of seconds. Abl2 FRAP was fit to a one-phase exponential equation that was modified from the above equation by excluding the second term.

### Spine head width measurement

Spine head width was measured in ImageJ using methods previously published with some modifications (Adrian, Kusters, Storm, Hoogenraad, & Kapitein, 2017; Nagerl, Willig, Hein, Hell, & Bonhoeffer, 2008). In short, spine head width was measured by drawing a line on the short axis of the spine. The fluorescence intensity profile was then plotted and fit to the following gaussian curve:

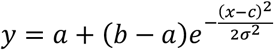

where *a* is the baseline intensity, *b* is the curve’s peak intensity, *c* is the position of the center of the peak and *σ* is the standard deviation. FWHM was then defined as 2.35 ∗ *σ*. For spine head width measurements from live samples the FWHM was averaged from all pre-bleached frames to account for local drifts of the spine in the focal plane.

### Experimental Design and Statistical Analysis

All data were from at least 3 independent neuronal cultures per condition. Imaging and analyses were performed while blinded to experimental conditions. Statistical analyses were performed in GraphPad Prism 8 or R software. Specific details on ‘n’ and statistical tests are included in the figure legends. Significance was defined by a p value less than 0.05 and a “*” symbol is always used for comparisons to the control group of the dataset. All pooled data are represented as mean ± SEM. Normality of data was checked with a D’Agostino and Pearson normality test where appropriate. In cases with simple genetic manipulations, an unpaired t-test was used. To analyze the effects of knockdown and rescue experiments where many manipulations were performed, we used one-way ANOVA to test for differences between groups. We performed a Pearson Correlation test to test for a relationship between cortactin spine enrichment and spine head width.

## Acknowledgements

We thank Aaron Levy, Josie Bircher, Melissa Carrizales and Amanda Jeng for helpful comments on this manuscript, as well as other members of the Koleske lab for their critical discussion and suggestions. This work was supported by NIH grants NS089662 (A.J.K.), NS105640 (A.J.K. and Michael J. Higley), MH115939 (A.J.K.), F31 MH116571 (J.E.S.), and J.E.S. was supported on NIH training grant T32GM007223 (Susan J. Baserga).

